# A SOSEKI-based coordinate system interprets global polarity cues in Arabidopsis

**DOI:** 10.1101/479113

**Authors:** Saiko Yoshida, Alja van der Schuren, Maritza van Dop, Luc van Galen, Shunsuke Saiga, Milad Adibi, Barbara Möller, Peter Marhavy, Richard Smith, Jiri Friml, Dolf Weijers

**Affiliations:** Laboratory of Biochemistry, Wageningen University, Stippeneng 4, Wageningen, the Netherlands; Institute of Science and Technology, Klosterneuburg, Austria; Department of Comparative Development and Genetics, Max Planck Institute for Plant Breeding Research, Cologne, Germany

## Abstract

Multicellular development requires coordinated cell polarization relative to body axes, and translation to oriented cell division. In plants, it is unknown how cell polarities are connected to organismal axes and translated to division. Here, we identify *Arabidopsis* SOSEKI (SOK) proteins that integrate apical-basal and radial organismal axes to localize to polar cell edges. Localization does not depend on tissue context, requires cell wall integrity and is defined by a transferrable, protein-specific motif. SOK proteins structurally resemble the DIX oligomerization domain in the animal Dishevelled polarity regulator. The DIX-like domain self-interacts and is required for edge localization and for influencing division orientation. Our work identifies a plant compass, interpreted by SOK proteins. Furthermore, despite fundamental differences, polarity in plants and animals converge upon the same protein domain.

---

Development of multicellular organisms relies on the ability to organize cell differentiation and division relative to body axes and to other cells within the same tissue. Individual cells are polarized, and polarity information is relayed to trigger local outgrowth^1,2^ or steer division orientation^1,3^. Mechanisms underlying polarization and division orientation are relatively well-understood in yeast and animals^4,5^. In plants, oriented division is critical for normal development^6,7^, and several polarly localized proteins have been identified^8–15^. However, yeast and animal polarity regulators seem to be missing from plant genomes, and it is thought that polarity components and mechanisms are distinct in plant and animal kingdoms^16^. Components of such mechanisms are elusive. The plant signaling molecule auxin regulates pattern formation, and defects in auxin response often manifest as changes in growth direction or cell division plane^6,17^. Mutations in the *Arabidopsis* MONOPTEROS/AUXIN RESPONSE FACTOR5 (MP) transcription factor^18^ cause alterations in division planes in the early embryo^19^. We argued that mediators of MP function in controlling cell division orientation should be among its transcriptional targets. Starting from a set of MP-dependent genes^20^, we here identify a family of novel, polarly localized proteins that link organismal axes to cell polarity and division orientation.

We previously performed transcriptome analysis on globular-stage embryos in which MP activity was locally inhibited^20^, and identified *TMO7* ^21^ as the most strongly down-regulated gene^20^. The second most strongly down-regulated gene (7.5-fold) is a gene of unknown function, containing a Domain of Unknown Function 966 (DUF966) and named *SOSEKI1* (explained below; *SOK1*; At1g05577). *SOK1* has 4 paralogues in the *Arabidopsis* genome: *SOK2* (At5g10150), *SOK3* (At2g28150), *SOK4* (At3g46110) and *SOK5* (At5g59790) (Fig. 1a; Extended Fig. 1) of which *SOK5* was also 2.4-fold down-regulated in embryos with reduced MP function.

**Fig 1:**
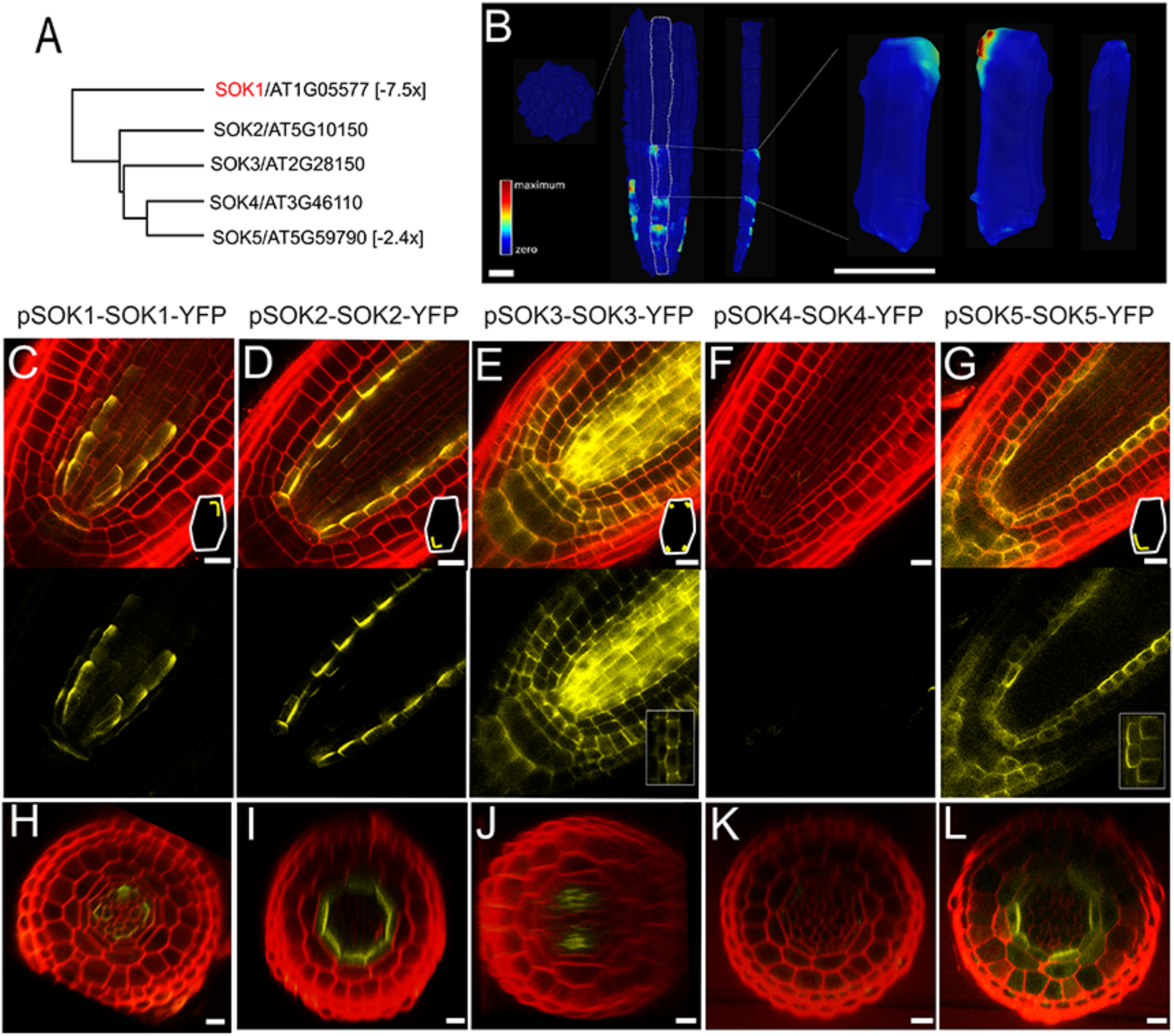
The SOSEKI family of polarly localized proteins. (a) Phylogenetic tree of the Arabidopsis DUF966/SOSEKI protein family. Values in brackets indicate fold-change (downregulation) in Q0990≫bdl embryos^5^. (b) SOK1-YFP protein localization in a root tip. Fluorescence values are shown in false color (red=maximum; blue=zero) on segmented cell surfaces. From left to right: cross-section of vascular cylinder, surface view of vascular cylinder, single cell file, and 3 different views of a single segmented cell. (c-l) Localization of SOK1-YFP (c,h), SOK2-YFP (d,i), SOK3-YFP (e,j), SOK4-YFP (f,k) and SOK5-YFP (g,l) in longitudinal cross-sections (c-g) and transverse cross-sections (h-l) of primary root meristems counterstained with Propidium Iodide (red). Insets in c,d,e,g schematically show subcellular SOK protein localization. Bars 10 µm.

To determine gene expression domains, nuclear-localized triple GFP (n3GFP) was driven by each *SOK* promoter, and this revealed that all *SOK* genes are expressed during embryogenesis and in primary/lateral roots (Extended Fig. 2). To observe protein localization, C-terminal YFP translational fusions were generated. All SOK-YFP patterns mirror pSOK-n3GFP patterns (Fig. 1, 2; Extended Fig. 2). Strikingly, each SOK protein marked novel cellular domains. SOK1-YFP marks the outer/apical edge of young vascular cells, including the pericycle, and in the columella root cap in the primary root (Fig. 1b,c,h). During embryogenesis, SOK1-YFP is first detected in the apical side of lower tier inner cells at the early globular stage (Fig. 2a). Subsequently, SOK1 localizes to the outer apical corner or outer lateral side of vascular cells and outer corners of the hypophysis at transition to heart stage (Fig. 2f; Extended Fig. 3). This pattern is maintained in the post-embryonic root (Fig. 1), and the lateral root (Fig. 2k; Extended Fig. 4). This localization pattern was confirmed by SOK1-tdTomato (Extended Fig. 2). The protein was named SOSEKI1 (Japanese for “cornerstone”) for this unique corner localization pattern. The SOK1 protein is highly unstable: SOK1-YFP signal disappears during cell division but is afterwards quickly re-established in lateral root primordia (Fig. 2p) and primary root (Fig. 2q). Treatment of SOK1-YFP roots with the translation inhibitor Cycloheximide (CHX) confirmed protein instability (Extended Fig. 6).

**Fig 2:**
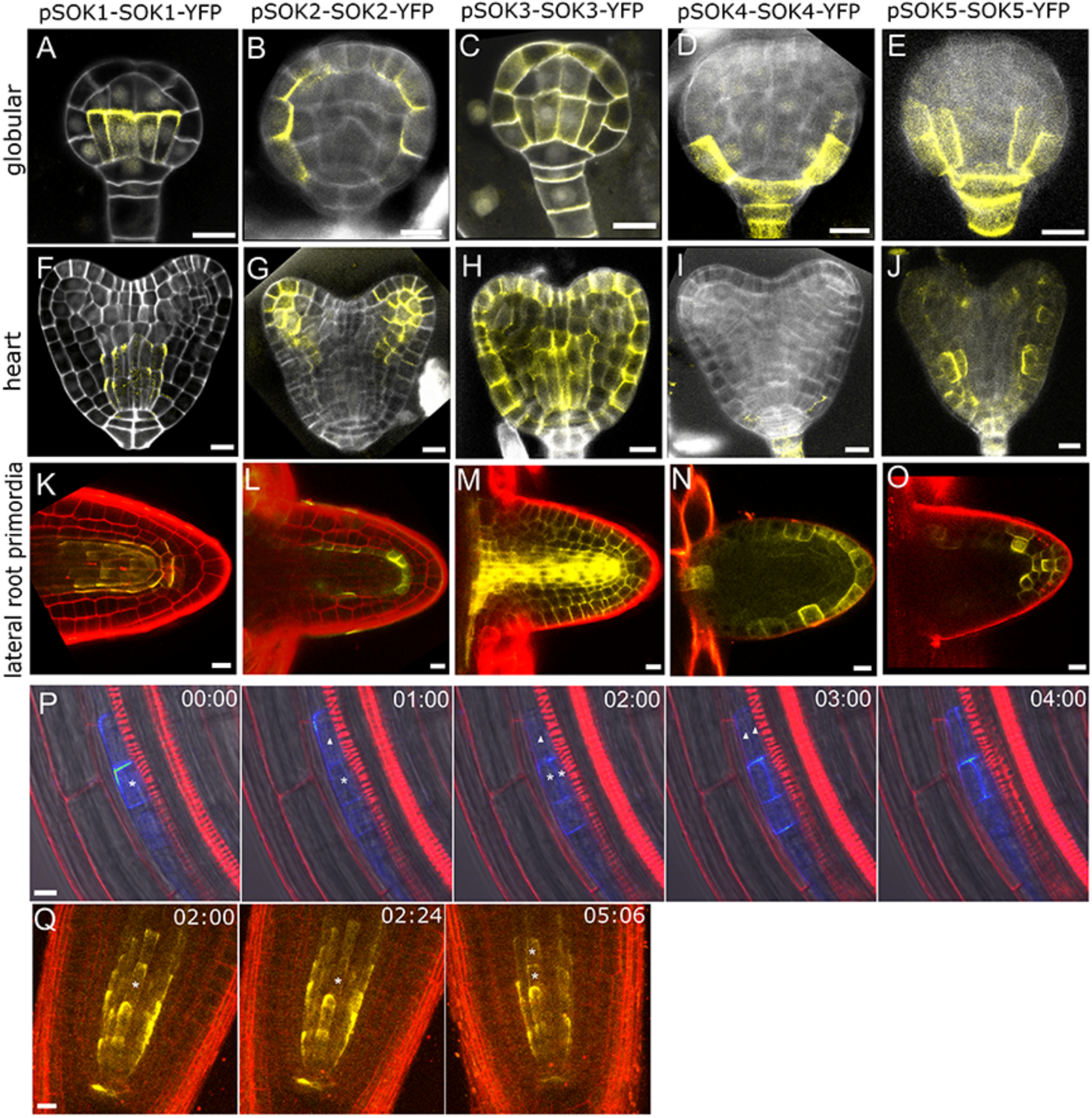
Diverse polar patterns of SOSEKI proteins. Localization of SOK1-YFP (a,f,k), SOK2-YFP (b,g,l), SOK3-YFP (c,h,m), SOK4-YFP (d,i,n) and SOK5-YFP (e,j,o) in globular stage embryos (a-e), heart stage embryos (f-j) and emerged lateral root primordia (k-o). (p,q) Stills of a time-lapse imaging series of SOK1-YFP fluorescence in an initiating lateral root (p) and in the primary root tip (q). Time (in hours) is indicated in each panel and dividing cell lineages are marked by asterisks. Embryos in a-j are counterstained with Renaissance RS2200 (white), and roots (k-q) with Propidium Iodide (red). Bars 10 µm.

All other SOK proteins were localized to different subcellular edges. SOK2-YFP localized to the inner basal edge of endodermal cells in the primary (Fig. 1d,i) and lateral root (Fig. 2l; Extended Fig. 4), starting in the globular embryo (Fig. 2b; Extended Fig. 3). SOK3-YFP accumulated at the basal side and all corners of most cells in the primary root (Fig. 1e,j), lateral roots (Fig. 2m; Extended Fig. 4) and embryos (Extended Fig. 3). Only faint SOK4-YFP accumulation was observed few vascular cells in the primary root (Fig. 1f,k), while it was expressed in lateral root (Fig. 2n; Extended Fig. 4) and embryo (Fig. 2d,i; Extended Fig. 3). SOK5-YFP resembles SOK2 localization in endodermis, QC and lateral root cap in primary root (Fig. 1g,l), yet its embryonic pattern evolves differently (Fig. 2e,j; Extended Fig. 3). Thus, all SOK proteins are expressed in specific tissues during root initiation, growth and branching, and mark unique polar subcellular domains.

To test whether differences in polar localization among SOK proteins are caused by cell types or protein determinants, we expressed SOK1-YFP or SOK2-YFP from the ubiquitous *RPS5A* promoter^22^. Apical SOK1 polarity was found in all cell types, but misexpressed SOK1 localizes to inner rather than outer edges in cortex and epidermis (Fig. 3a-c). Ectopic SOK2-YFP localizes to inner basal edges of all cells (Fig. 3e). Thus apical/basal polarity is maintained in misexpression lines and appears intrinsic to SOK1 and SOK2 proteins. Strikingly, in these lines, polar localization was found across the embryonic shoot/root axis (Fig. 3d,f), suggesting the existence of a common polarity reference in the entire body. During lateral root initiation however, localization followed the new organ axis (Fig. 2k,l; Extended Fig. 4), suggesting that the SOK-based coordinate system is autonomous to lateral organs. In contrast to apical/basal polarity, inner/outer polarity of SOK1 depended on position relative to the endodermis (Fig. 3a); SOK1 always localized pointing towards the endodermis. Localization in the *shortroot* (*shr*) and *scarecrow* (*scr*) mutants with impaired endodermal identity^23,24^ caused loss of edge localization and led to apical accumulation in the mutant cell file (Fig. 3k,l). This suggests that edge localization integrates genetically separable apical-basal and outer-inner axes. The cortex-endodermis junction serves as a potent cue for SOK1 localization, which is confirmed by ground tissue-specific expression of SOK1-YFP using the N9135 GAL4 driver line^25^. In the shared initial for cortex/endodermis, SOK1-YFP is apical, while the protein localized at both opposing edges toward the junction after the periclinal division separating separates endodermis and cortex (Fig. 3m,n). These observations revealed that plant cells and tissues possess a universal coordinate system with an internal reference that is read and integrated by SOK proteins.

**Fig 3:**
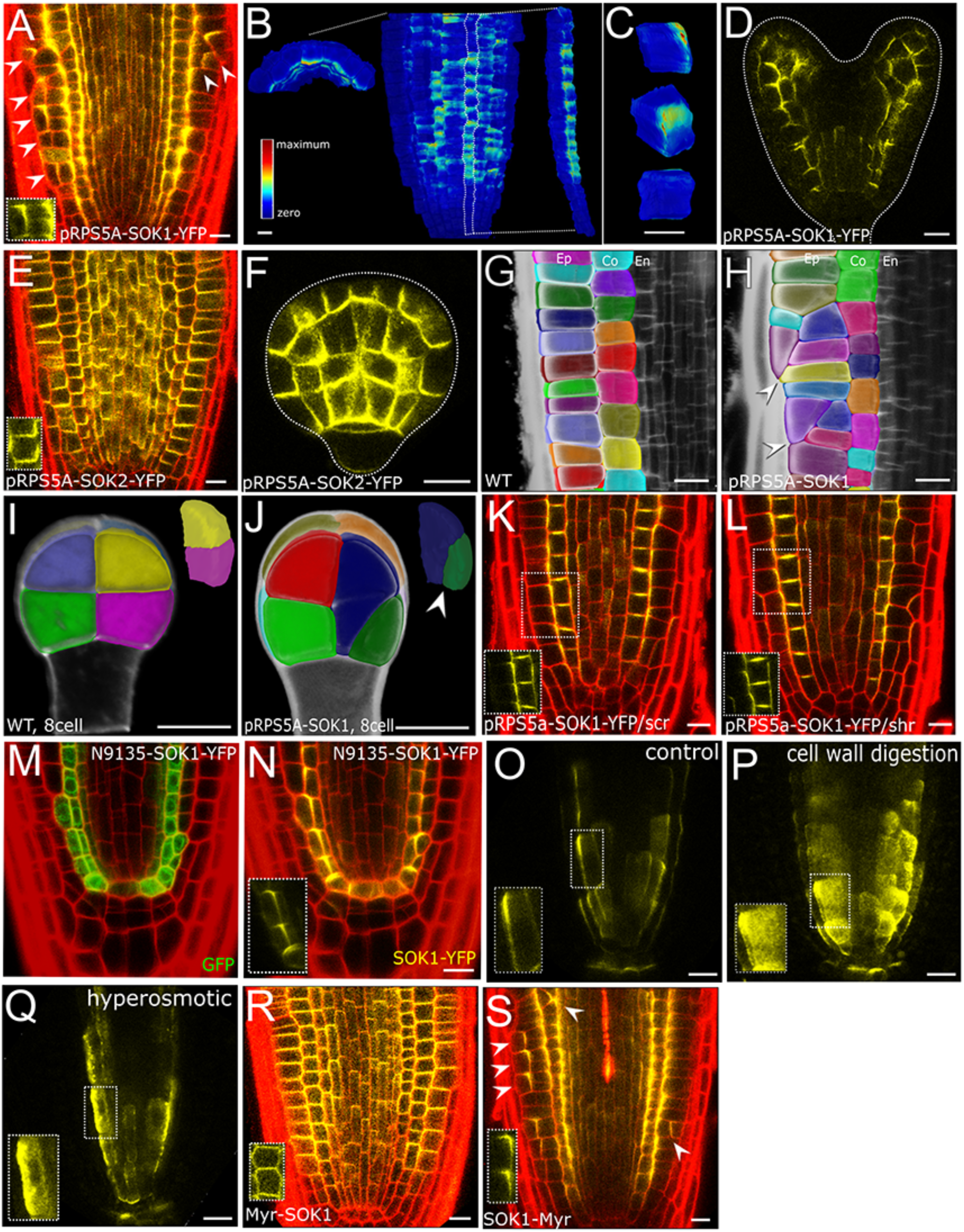
Regulation of SOSEKI localization. (a) Localization of SOK1-YFP in RPS5A-SOK1-YFP root tip. Arrowheads indicate oblique cell divisions. (b,c) SOK1-YFP protein localization in a RPS5A-SOK1-YFP root tip (b) and 3D localization of SOK1-YFP on segmented epidermal cells (c). Fluorescence values are shown in false color (red=maximum; blue=zero) on segmented cell surfaces. (d) SOK1-YFP localization in RPS5A-SOK1-YFP heart stage embryo. (e,f) Localization of SOK2-YFP in RPS5A-SOK2-YFP root tip (e) and globular stage embryo (f). (g-j) Segmented cell volumes in wild-type (g,i) and RPS5A-SOK1 (h,j) root meristem (cells located between 70-160 μm away from the quiescent center were observed in g,h) and 8-cell embryo (i,j). (k,l) Localization of *RPS5A*-driven SOK1-YFP in *scr* ((k) and *shr* (l) root tips. (m,n) GAL4-driven GFP expression (m) and GAL4-driven SOK1-YFP accumulation (n) in root tip of N9135≫SOK1-YFP. (o-q) Localization of SOK1-YFP in control-treated root tip (o), and in root tips treated with cell wall-digesting enzymes for 3 min (p) or with 0.4 M mannitol for 1.5 h (q). (r) Myr-SOK1-YFP localization in RPS5A-Myr-SOK1-YFP root tip. (s) SOK1-Myr-YFP localization in RPS5A-SOK1-Myr-YFP root tip. Arrowheads indicate oblique cell divisions. Walls in (a,e,g-n,r,s) are counterstained with Propidium Iodide (red). Insets in (a,e,k-n,r,s) show SOK protein localization in single cells. Bars 10 µm.

Polar localization of plant proteins generally depends on dynamic processes such as membrane trafficking, cytoskeletal dynamics or protein degradation^8–10,12,14^. In stark contrast, none of 20 tested hormone or drugs affecting these processes, nor changes in light, temperature or gravity angle affected the edge localization of SOK1, SOK2 or SOK3 (Extended Fig. 6; Extended Table 1). This suggests that polar localization follows a pathway that is different from well-known polar proteins such as PINs, BOR1, NIP5 and PEN3 ^8–10,12,14^. We next tested if the cell wall, or mechanical properties influence SOK localization and found that a brief (<20 minutes) treatment with cell wall-degrading enzymes or high osmotic mannitol solution led to internalization and mis-localization of SOK proteins (Fig. 30-q; Extended Fig. 5). This indicates that cell wall integrity is critical for edge localization and membrane association of SOK proteins.

SOK1-YFP misexpression induced frequent alterations in cell division orientation; in embryos, such divisions were found in all cell types (Fig. 3i,j; Extended Fig. 7), while in roots, defects were strongest in the cortex and epidermis (Fig. 3a,g,h). While root cells normally divide anticlinal to the growth axis (Fig. 3g), misexpression lines displayed either oblique or periclinal divisions generating additional cell layers (Fig. 3h). The same defect was induced by misexpressing non-tagged SOK1, and was accompanied by slightly inhibited root growth (Extended Fig. 7). We utilized this activity in redirecting cell division planes to determine protein determinants and properties required for activity. SOK proteins do not have a signal peptide or predicted transmembrane helices (Extended Fig. 1) and are likely peripherally membrane-associated. We fused a Myristoylation (Myr) motif ^26^ to either the N-or C-terminus of SOK1-YFP. Both polarity and activity (as judged by oblique cell divisions) were completely lost when the Myr motif was fused to the N-terminus of SOK1-YFP (Fig. 3s), while adding the Myr motif to the C-terminus of SOK1-YFP did not affect either (Fig. 3r). Thus, insertion in the plasma membrane prevents polar localization, which is in turn required for SOK1 function.

To locate polarity determinants, we generated a series of N-or C-terminal deletions (Fig. 4a-e; Extended Fig. 8) and misexpressed each as a YFP fusion. Deletions ΔA, ΔB, ΔC and ΔD caused SOK1 to be localized in the cytosol. Interestingly, ΔE localized to the apical edge, suggesting that the fragment contained in the ΔD-ΔE segment is sufficient for polar localization and for altering division orientation. Deletions ΔF, ΔG, ΔH and ΔI are broadly localized to plasma membrane, suggesting that the N-terminus is not required for localized membrane association per se, but rather for focusing to the edge. Deletions ΔJ, ΔK and ΔL were all localized to the cytosol, suggesting that the ΔD-ΔE segment can only direct edge localization if the N-terminus is present. Thus, SOK1 carries two function domains: one for membrane association (middle), and another (N-terminal) for focused polar localization. Both are required for activity in changing division orientation.

**Figure 4:**
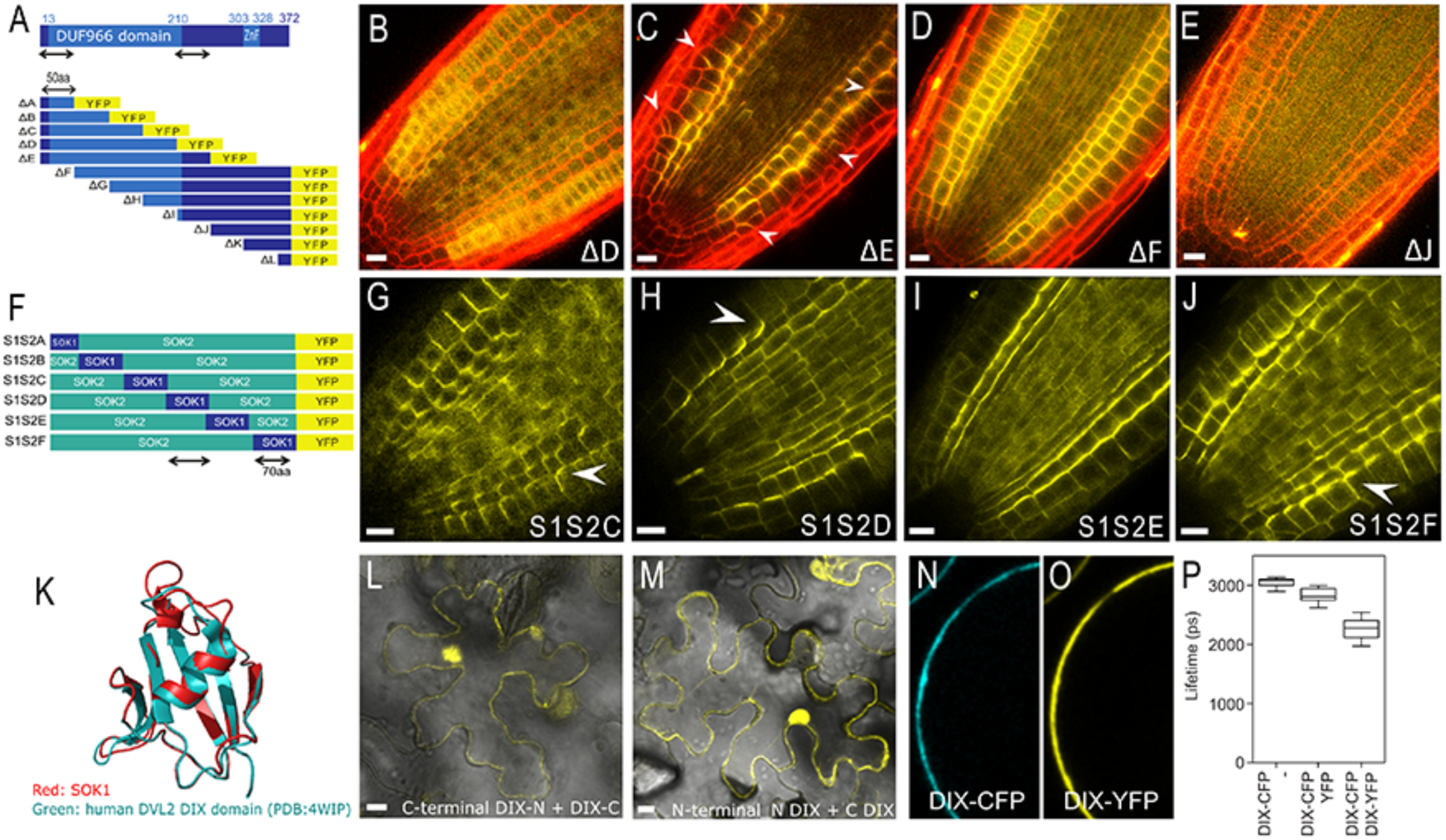
Protein determinants for SOK localization and activity. (a) Schematic of SOK1 protein domains (ZnF=Zn-finger), and outline of domain deletion constructs expressed as YFP-fusions driven from the *RPS5A* promoter. Numbers indicate amino acid positions relative to the start. (b-e) Representative examples of SOK1-YFP domain deletion accumulation in root tips. Arrowheads in (c) indicate oblique divisions. (f) Outline of SOK2-SOK1 domain swaps, expressed as YFP fusions driven from the *RPS5A* promoter. (g-j) Representative examples of SOK1-SOK2-YFP domain swap accumulation in root tips. Arrowheads in (g) and (j) mark basal localization, while arrowhead in (h) marks apical localization. Note that regions in (a) and (f) marked by arrows denote the regions for focused membrane localization and polarity defined by deletions and swaps. (k) Structural alignment of the DIX domain in human DVL2 (PDB: 4WIP; blue) and the homology model of the SOK1 DIX-like domain (red). (l,m) Bimolecular Fluorescence complementation in *Nicotiana benthamiana* epidermis of the SOK1 DIX-like domain in homo-dimeric conformation, using either N-terminal YFP fragments (l) or C-terminal YFP fragments (m). (n,o) Fluorescence of co-expressed SOK1 DIX-like domain as CFP (n) and YFP (o) fusion in protoplast. (p) Quantification of CFP Fluorescence Lifetime (in picoseconds, ps) in SOK1-DIX-CFP/YFP pair, and the two individual fusions in protoplasts (n=20 protoplasts for each fusions). Walls in (b-e) are counterstained with Propidium Iodide (red). Bars 10 µm.

To test whether different SOK proteins use similar domains for localization, we replaced successive 50-70 amino acids of the basally localized SOK2-YFP by the corresponding region of SOK1 (Named S2S1A-F; Fig. 4f-j; Extended Fig. 8). While most chimaeras localized to the basal edge (Fig. 4g,j; Extended Fig. 8), S2S1D shifted to the apical edge (Fig. 4h; Extended Fig. 8), similar to SOK1-YFP. The S1S2E showed a mixture of SOK1 and SOK2 localization, at the inner lateral membrane (Fig. 4i; Extended Fig. 8). Thus, polar localization can be transferred between SOK1 and SOK2 using a discrete domain that overlaps with the ΔD-ΔE “polarity domain” defined by deletions. We confirmed that the same domain conferred polarity changes in swaps between SOK1 and SOK5 (Extended Fig. 9). This identifies a minimal domain for polar targeting of multiple SOK members.

SOK proteins are plant-specific, and the function of DUF966 domain is unknown. We used structural homology modeling to identify potential homologues, and found that, despite very low sequence conservation, a part of the DUF966 domain resembles the DIX (Dishevelled/Axin) domain (Fig. 4k) that is found in a number of animal proteins^27,28^. The DIX domain mediates head-to-tail oligomerization^29^ and is required for clustering of the polarity regulator Dishevelled in *Drosophila*^28,29^. Bimolecular Fluorescence Complementation in tobacco leaves (Fig. 4l,m; Extended Fig. 10) and a FRET assay in *Arabidopsis* protoplasts (Fig. 4n-p) showed that the predicted DIX domain in SOK1 indeed homodimerizes. Importantly, the DIX-like domain corresponds to the N-terminal region that is required for edge localization and biological activity (Fig. 4a,d; Extended Fig. 1). Thus, the animal cell polarity protein Dishevelled and SOSEKI1 may use the same protein domain for their asymmetric localization and polarity-related function.

Our study identified a novel plant-specific family of polar, edge-localized proteins. Localization to previously unidentified polar domains is important for the activity of SOK1 in influencing the cell division axis. Whether SOK1 and other family members mediate this function, and what cellular mechanism underlies such activity is yet to be determined. Nevertheless, our analysis of SOK localization identified a universal coordinate system in plants that extends well beyond the global polarity field in the leaf^30^, and encompasses the entire body axis. SOK proteins that can locally interpret and integrate the global coordinates. Consistent with being the output of a coordinate system, SOK localization is robust, yet seems to rely on cell wall integrity and/or mechanical properties. Further exploration of the mechanisms of SOK localization and function may help revealing the fundamental principles of cell and organismal polarity in plants. This may herald surprising analogies to animal polarity mechanisms, given the adoption of the same functional protein domain in polar proteins across kingdoms.

## Acknowledgments

We thank Dominique Hagemans, Vivienne Mol and Colette ten Hove for support and advice with experiments, and Eva Benkova for supporting P.M.. This work was supported by the European Research Council under the European Union’s Seventh Framework Programme (FP7/2007-2013) under REA grant agreement n° [291734], and a Marie Curie Fellowships (contract 753138) to S.Y., by a European Research Council grant (ERC-StG CELLPATTERN; contract 281573), and ALW Open Competition grant (820.02.019) and an ALW-VIDI grant (864.06.012) from the Netherlands Organization for Scientific Research (NWO) to D.W., a grant (831.13.001) from the Netherlands Organization for Scientific Research (NWO) to M.v.D., and a FEBS long-term fellowship to P.M..

## Author contributions

D.W. conceived the study. S.Y. performed most of the experiments under the supervision of D.W., and with help from A.v.d.S., L.v.G. and S.S.. M.v.D. Performed BiFC and FRET-FLIM experiments, SOK1 localization in ground tissue and in *shr* and *scr* mutants, and SOK1/SOK5 swaps. B.M. initiated the project and identified the *SOK1* gene. M.A. and R.S.S. performed 3D SOK1 localization analysis. P.M. helped to perform live imaging under J.F.’s supervision. S.Y. and D.W. wrote the manuscript with input from all authors.

