## Supplementary material for "A SOSEKI-based coordinate system interprets global polarity cues in Arabidopsis"

**This PDF file includes:**

Materials and Methods
Extended Figures 1-10
Extended Tables 1-2
Captions for Supplemental Movies 1-3

**Other Supplementary Materials for this manuscript include the following:**

Supplemental Movies 1-3

### Materials and Methods

Plant material

Arabidopsis ecotype Columbia-0 was used in all experiments except when noted otherwise. The N9135-GAL4 enhancer trap line[1] was generated by Jim Haseloff (University of Cambridge, UK) and obtained though the Arabidopsis Biological Resource Centre (ABRC). *shr*-2 and *scr*-4 mutants were previously described[2,3].

All seeds were sterilized, sown on MS medium with 1 % sucrose and 0.8% Daishin agar (Duchefa) and vernalized for one day. Plants were grown on soil at 22°C under the long-day condition.

Molecular cloning

All cloning was performed using previously described Ligation-Independent Cloning methods and vectors[4].For promoter-GFP lines, 1.2-5kb fragments upstream of the ATG of *SOK* genes were amplified and introduced into upstream of SV40-n3eGFP in pPLV04. For C-terminal translational fusion lines, genomic fragments, including 1.2-5 kb upstream of *SOK* genes, were introduced into pPLV16 (sYPF2) or pPLV22 (tdTomato). For misexpression lines, *SOK* cDNAs were amplified, fused with sYFP2 and introduced into pPLV28 downstream of the *RPS5A* promoter. For gene swap experiments, cDNA fragments of *SOK1*, *SOK2, SOK5* and YFP were amplified, fused by overlap extension PCR and introduced downstream of the *RPS5A* promoter in pPLV28. For deletion experiments, cDNA fragments from SOK1-YFP were amplified and introduced downstream of the *RPS5A* promoter in pPLV28. For myristoylated SOK1 constructs, sequences for myristoylation (GGCFSKK)[5] was added to either C-terminus or N-terminus of SOK1 cDNA sequences, and introduced downstream of the *RPS5A* promoter in pPLV28. The *UAS*::SOK1-YFP construct was generated by amplifying SOK1-YFP from the pRPS5A::SOK1-YFP plasmid and introducing it downstream of the GAL4-dependent *UAS* promoter [6] in pPLV32. The construct was transformed directly into the N9135 enhancer trap line[1]. For BiFC, the DIX cDNA sequence was amplified from the pRPS5A::SOK1-YFP plasmid and cloned into a modified pGreenII vector containing p35S::LIC-n/cYFP. FLIM vectors were generated by cloning the DIX cDNA into pMON 35S::LIC-sYFP or pMON 35S::LIC-sCFP3a.

All constructs were sequenced, and transformed into wild-type Columbia or N9135 GAL4 by floral dip using Agrobacterium strain GV3101(pSoup). All primers used for cloning are listed in the Extrended Table 2.

Microscopic analysis

Roots were stained by propidium iodide (PI; Sigma-Aldrich) at final concentration of 10µg/ml. Embryos were fixed and stained by 2.2% Renaissance RS2200 (Renaissance Chemicals Ltd) in PBS buffer at pH 6.9 containing 4% paraformaldehyde, 5% glycerol, 4.2% dimethyl sulfoxide (DMSO) or stained by PI after fixation as previously described[7,8].

Confocal imaging was performed with Zeiss LSM700/800 (observation of lateral root) or Leica SP5/II (observation of primary root and embryo) as previously described[8]. Live imaging was performed with LSM700 as described before[9,10]. For the 3D imaging of pSOK1-SOK1-YFP and pSOK1-GFP root, Zeiss LSM780 with 2 photon laser (960-990nm) and 500-550nm band-pass filter were used. Analysis of confocal images (3D reconstruction, 3D segmentation) was performed by MorphoGraphX [11].

Chemical treatments

All chemicals used to treat SOK-YFP roots are shown in Extended Table 1. Concentration for each chemical was obtained from literature. Seedlings were placed on MS plates containing each chemicals for the durations mentioned in the text and were subsequently imaged by confocal microscopy. As controls for the chemical treatments, the same percentage of DMSO or ethanol used for each chemical treatment were used and no effect was observed. For most of the chemical treatments, final concentrations of DMSO were <0.5% (v/v) and ethanol were <0.1% (v/v).

Plasmolysis was performed either on MS medium with mannitol (Sigma, M9546) at final concentration 0.4 M or by dipping roots in 0.4 M mannitol solution in milliQ water, and stained by FM4-64 at least for 2 min. For cell wall digestion, either 1% Cellulose R10 (Yakult Honsha Co. L.T.D, Japan) and/or 0.2% Macerozyme (Duchefa) were used. Cellulose and/or Macerozyme was dissolved in a solution containing 0.4 M mannitol, 10mM $CaCl_2$, 20 mM KCl and 20 mM MES (pH 5.7).

BiFC and FRET-FLIM

BiFC was performed as follows: *Agrobacterium* containing BiFC plasmids were grown overnight in 5 ml LB + 20 mg/L Gentamycin, 50 mg/L kanamycin, 25 mg/L rifampicin and 2 mg/L tetracyclin. Cultures were spun down at 4000 rpm for 10 minutes and the bacterial pellet was resuspended in 1 mL MMAi (5g/L MS salts without vitamins, 2g/L MES, 20g/L sucrose, pH 5.6; and 0.2 mM Acetosyringone). The $OD_{600}$ was measured with a spectrophotometer. The infiltration samples were mixed 1:1 at a total $OD_{600}$ of 0.8. Samples were incubated at RT for 2 hours and infiltrated into the underside of *Nicotiana benthamiana* leaves with a 1 mL syringe. After two days, leaf samples were cut out with a razor blade and imaged with a confocal microscope. Protoplast transfection and FLIM measurements were performed on a Leica SP8 as described[12].

Structural homology modeling

SwissModel (https://swissmodel.expasy.org/) was used to model the structure of DIX-LIKE. To this end, the conserved N-terminal part of SOK1 was entered into the program and the software itself selected the best matching crystal structure, which was human Dvl2 (PBD: 4WIP).

Statistics and image editing

For all experiments involving transgenic lines, care was taken to analyze multiple independent transgenic lines (n>2 when studying T2 or T3 generation; n>10 for T1 generation), where lines were selected based on representing non-extreme characteristics. For all microscopy analyses, at least 5 individuals were analyzed for each genotype/treatment. Only results representing the majority of observations are shown.

Raw microscopy images were collected, and brightness was uniformly modified across the entire image. When images were to be directly compared, brightness modifications were performed in an identical manner.

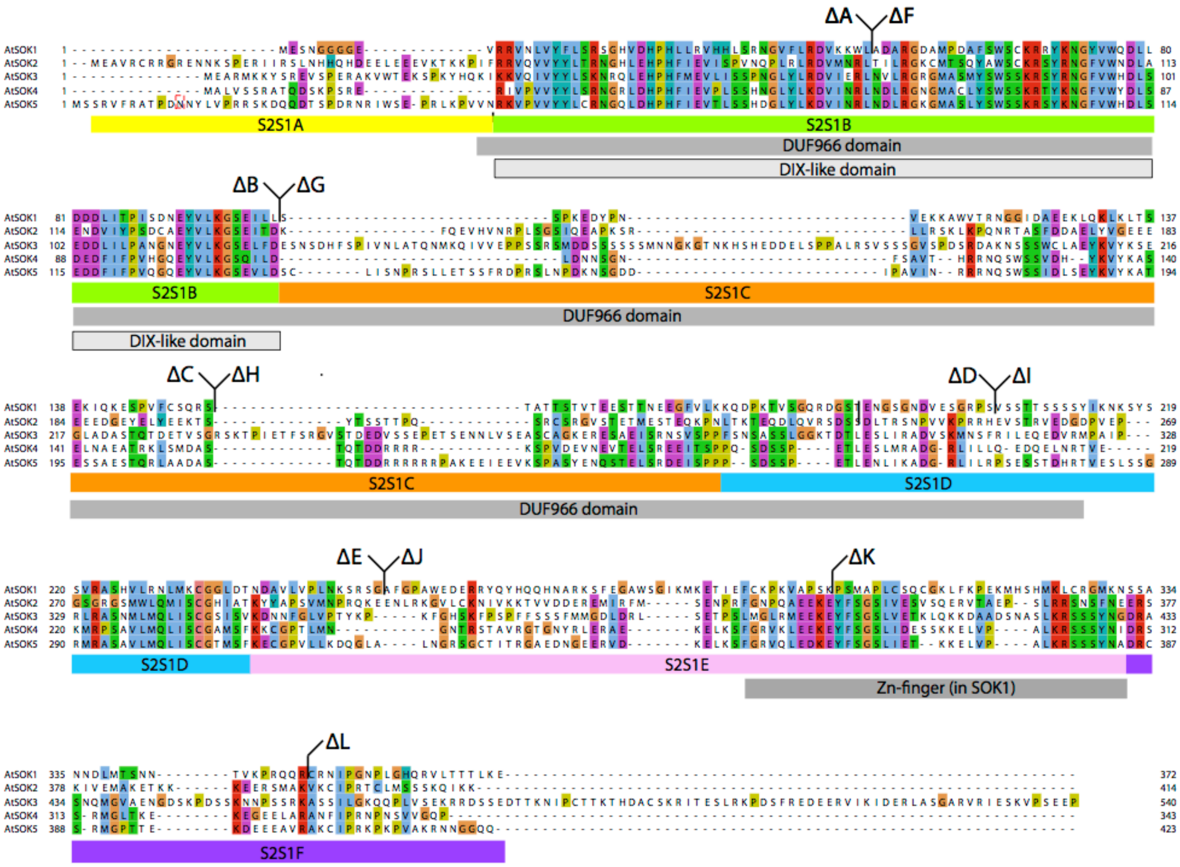


**Extended Figure 1: SOK protein alignment, domains, swaps and deletion fragments**
Protein sequence alignment of the 5 Arabidopsis SOK proteins. The DUF966 domain, as well as the DIX-like domain and the Zinc Finger are indicated. All deletion fragment boundaries are bracketed above the sequence, and the regions swapped between SOK1 and SOK2 are marked underneath the alignment.

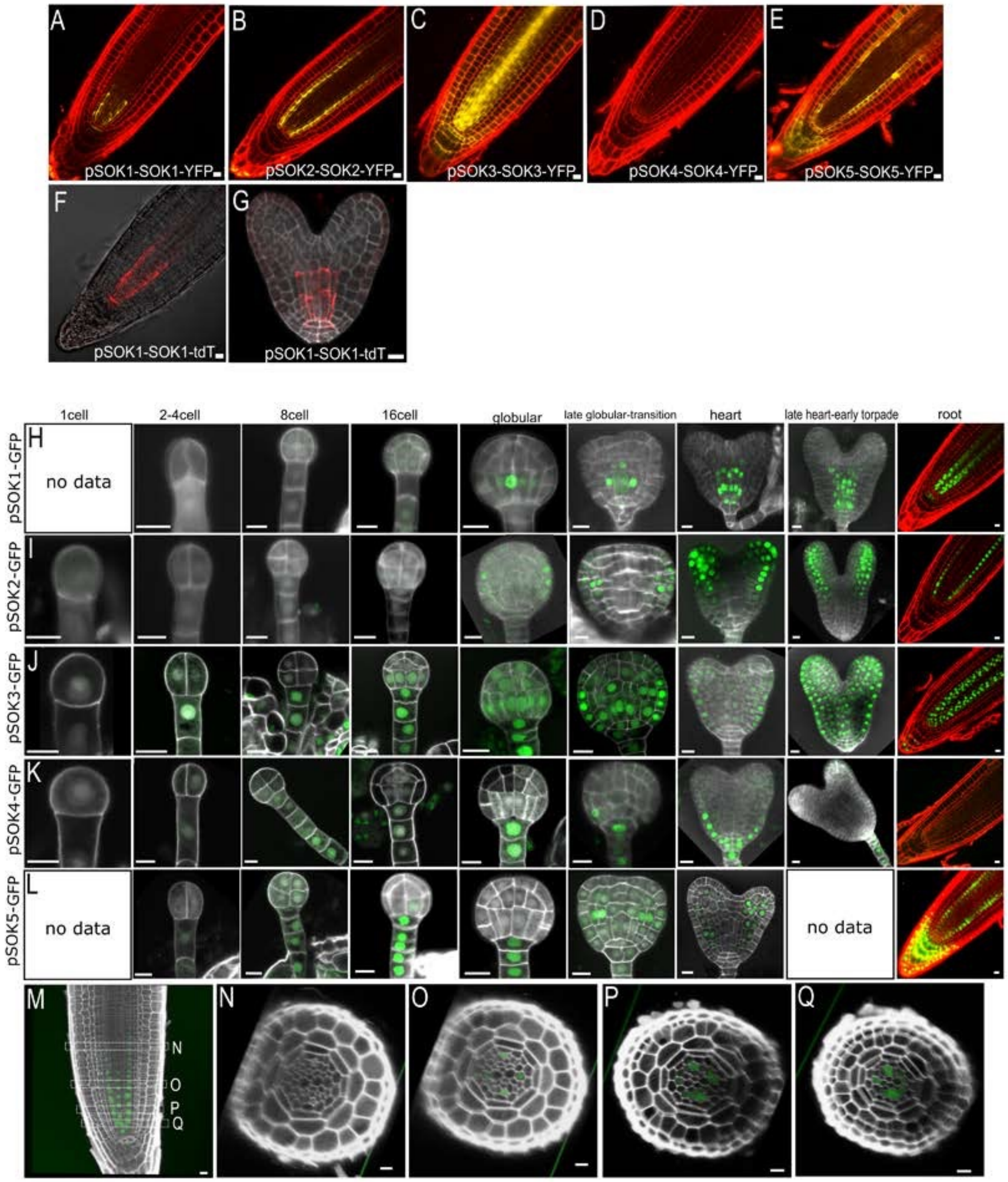


**Extended Figure 2: SOK gene expression and protein localization in embryo and root**

(a-e) Fluorescence of SOK-YFP fusion proteins in the extended root meristem, when expressed from the native promoter. (f,g) Fluorescence of SOK1-tdTomato in root tip (f) and heart stage embryo (g).

(h-l) Fluorescence of n3GFP in successive stages of embryo development and the root meristem of lines expressing *pSOK1*-n3GFP, *pSOK2*-n3GFP, *pSOK3*-n3GFP, *pSOK4*-n3GFP and *pSOK5*-n3GFP. (m-q) Longitudinal (m) and transverse (n-q) cross-sections through root expressing *pSOK1*-n3GFP. Embryos (g-l) are counterstained with Renaissance RS2200 (white), and roots (a-e, h-q) with Propidium Iodide. Bars 10 μm.

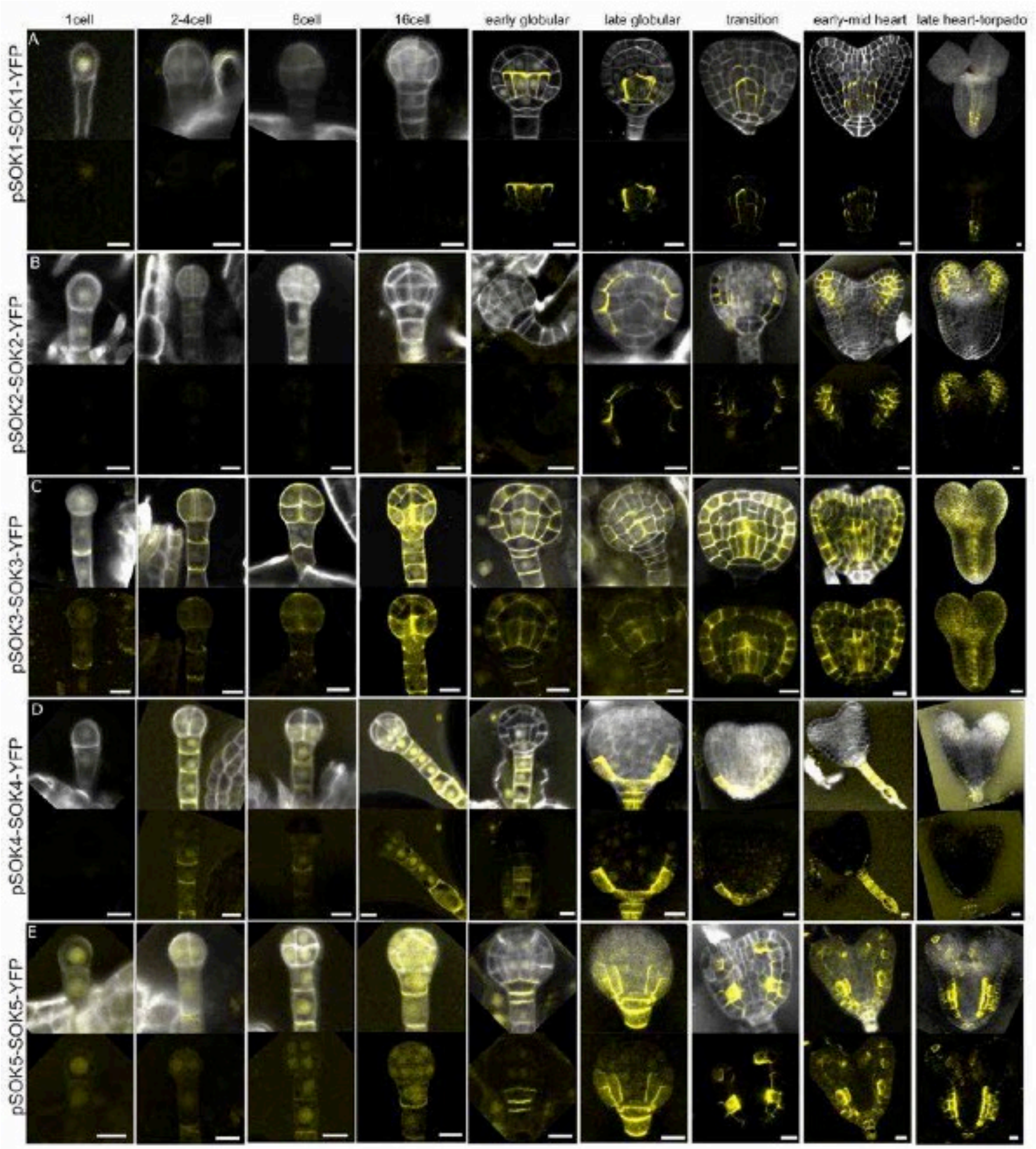


**Extended Figure 3: SOK protein localization during embryogenesis.**
SOK1-YFP (a), SOK2-YFP (b), SOK3-YFP (c), SOK4-YFP (d) and SOK5-YFP (e) protein accumulation during successive stages of embryogenesis. Upper panels show overlay of YFP and cell wall (Renaissance RS2200 – white) signals, while lower panels show only the YFP signal. Bars 10 µm.

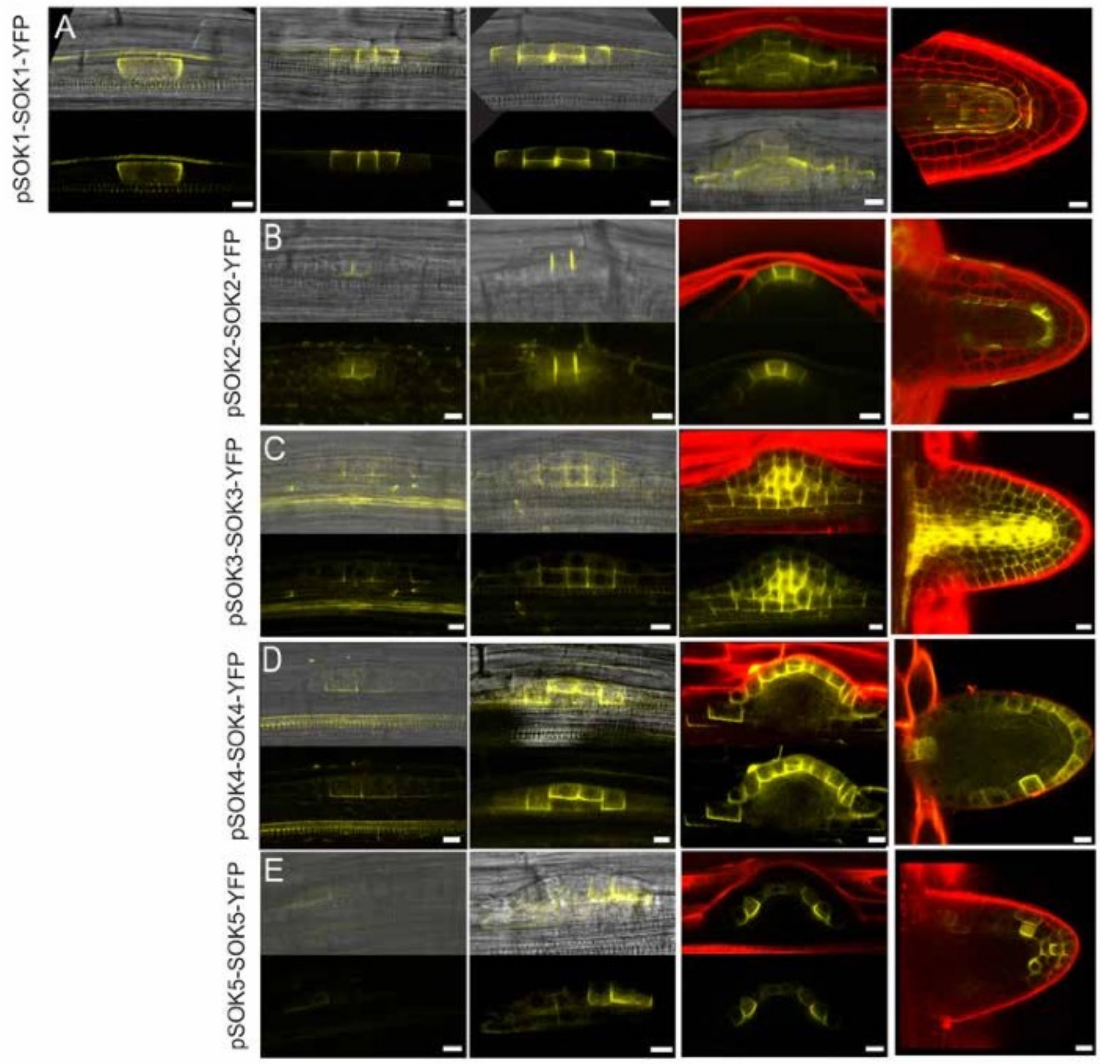


**Extended Figure 4: SOK protein accumulation during lateral root development.**
SOK1-YFP (a-e), SOK2-YFP (f-i), SOK3-YFP (j-m), SOK4-YFP (n-q) and SOK5-YFP (r-u) protein accumulation during successive stages of lateral root formation. Upper panels in (a-d,f-h,j-l,n-p,r-t) are overlays of YFP signal and transmitted light images (a-c,f,g,j,k,n,o,r,s) or Propidium Iodide (d,h,l,p,t - red), while lower panels are YFP signal alone. Bars 10 µm.

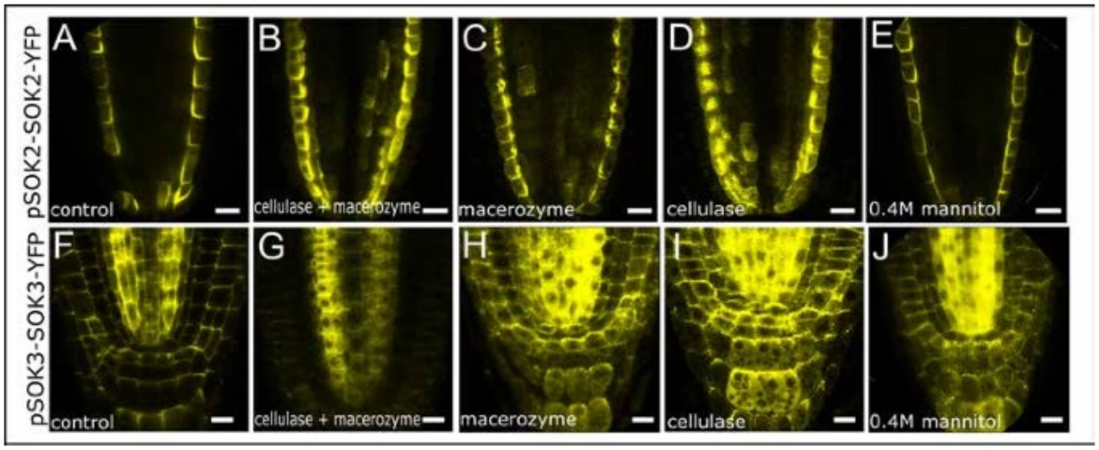


**Extended Figure 5: Cell wall-dependent SOK protein localization**
(a-j) Localization of SOK2-YFP (a-e) and SOK3-YFP (f-j) protein in control root (a,f) and root tips treated with Cellulase and Macerozyme (b,g), Macerozyme (c,h), Cellulase (d,i) or 0.4 M Mannitol (e,j). Bars 10 µm.

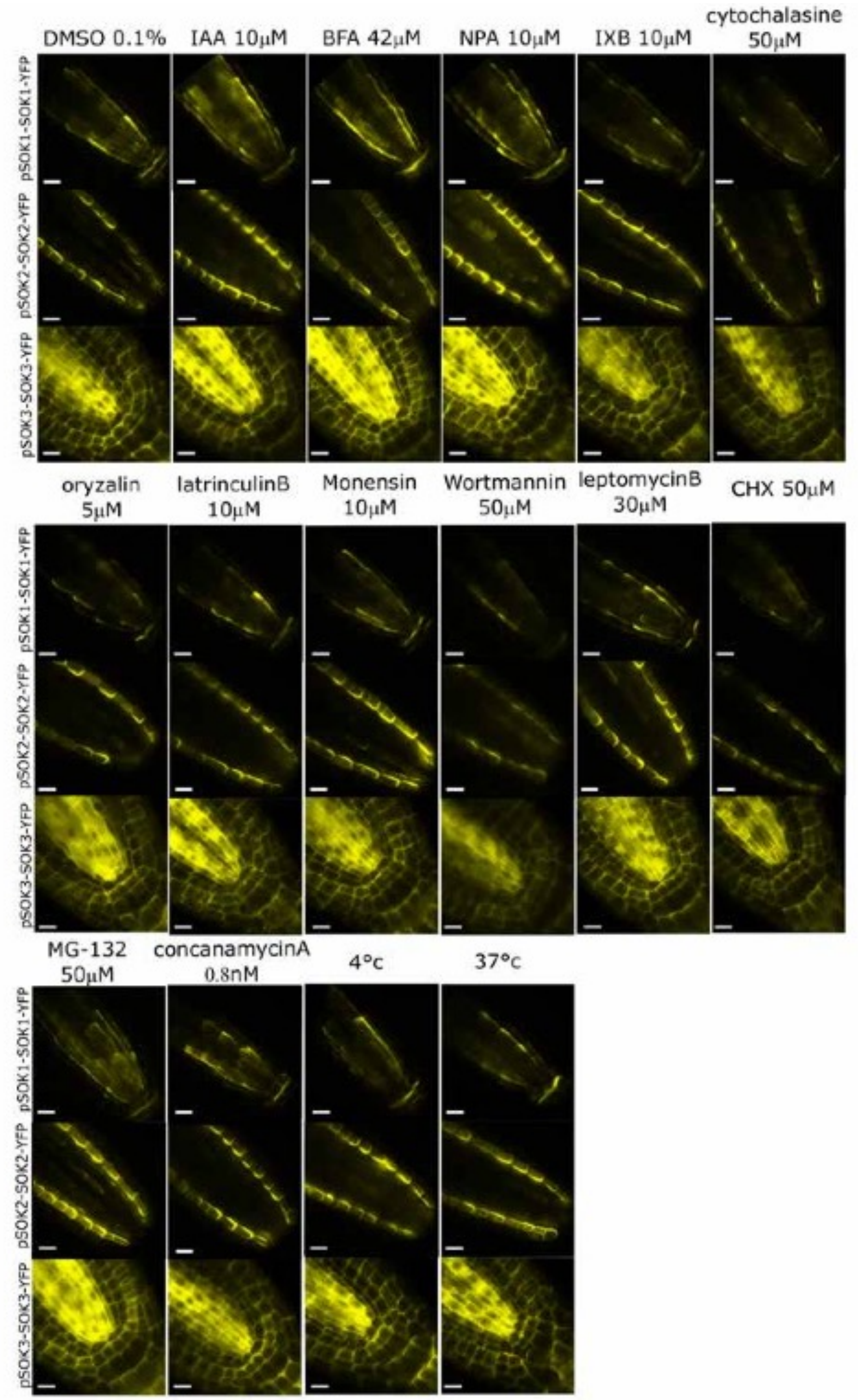


**Extended Figure 6: Pharmacological analysis of SOK protein localization requirements**
Localization of SOK1-YFP, SOK2-YFP and SOK3-YFP protein in control root (0.1% DMSO), and roots treated with a range of drugs and conditions as indicated above each column. Bars 10 µm.

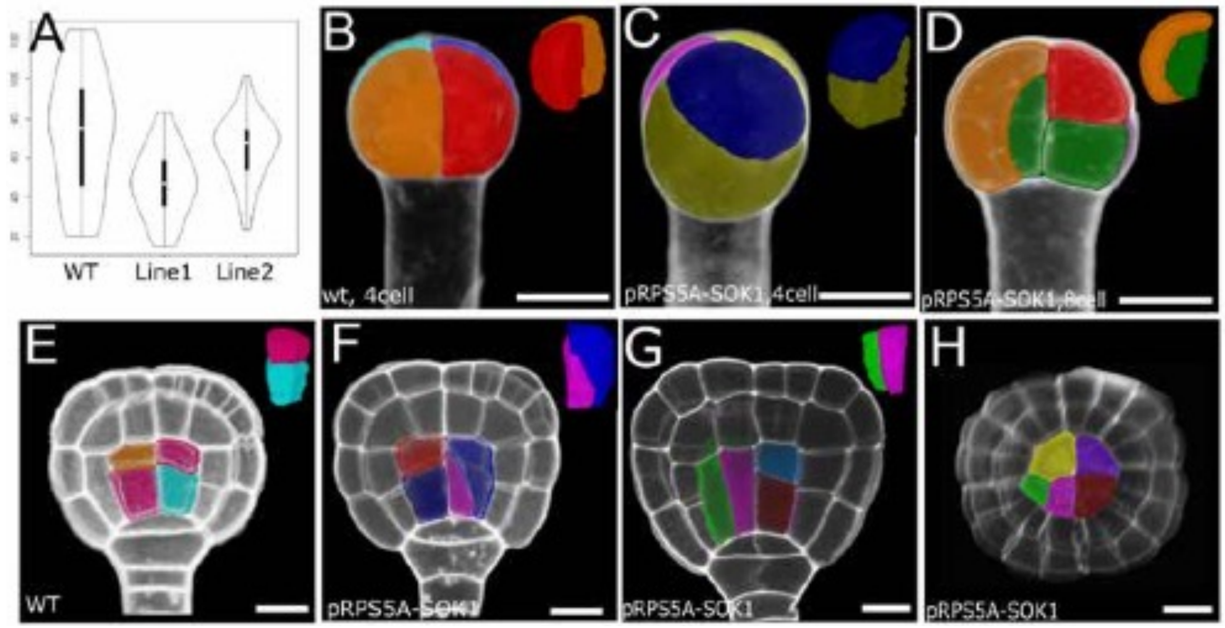


**Extended Figure 7: Defects induced by SOK1 misexpression**
(a) Root length (mm) in wild-type and two independent homozygous RPS5A-SOK1 lines, n=50 (wt), 44 (line1), 39 (line2). (b-h) Segmented cell volumes in wild-type (b,e) and RPS5A-SOK1 (c,d,f-h) embryos. Cell clusters, extracted from each embryo are shown in the top right corner of each panel. Note that (h) is a cross-section of the embryo shown in (g). Bars 10 µm.

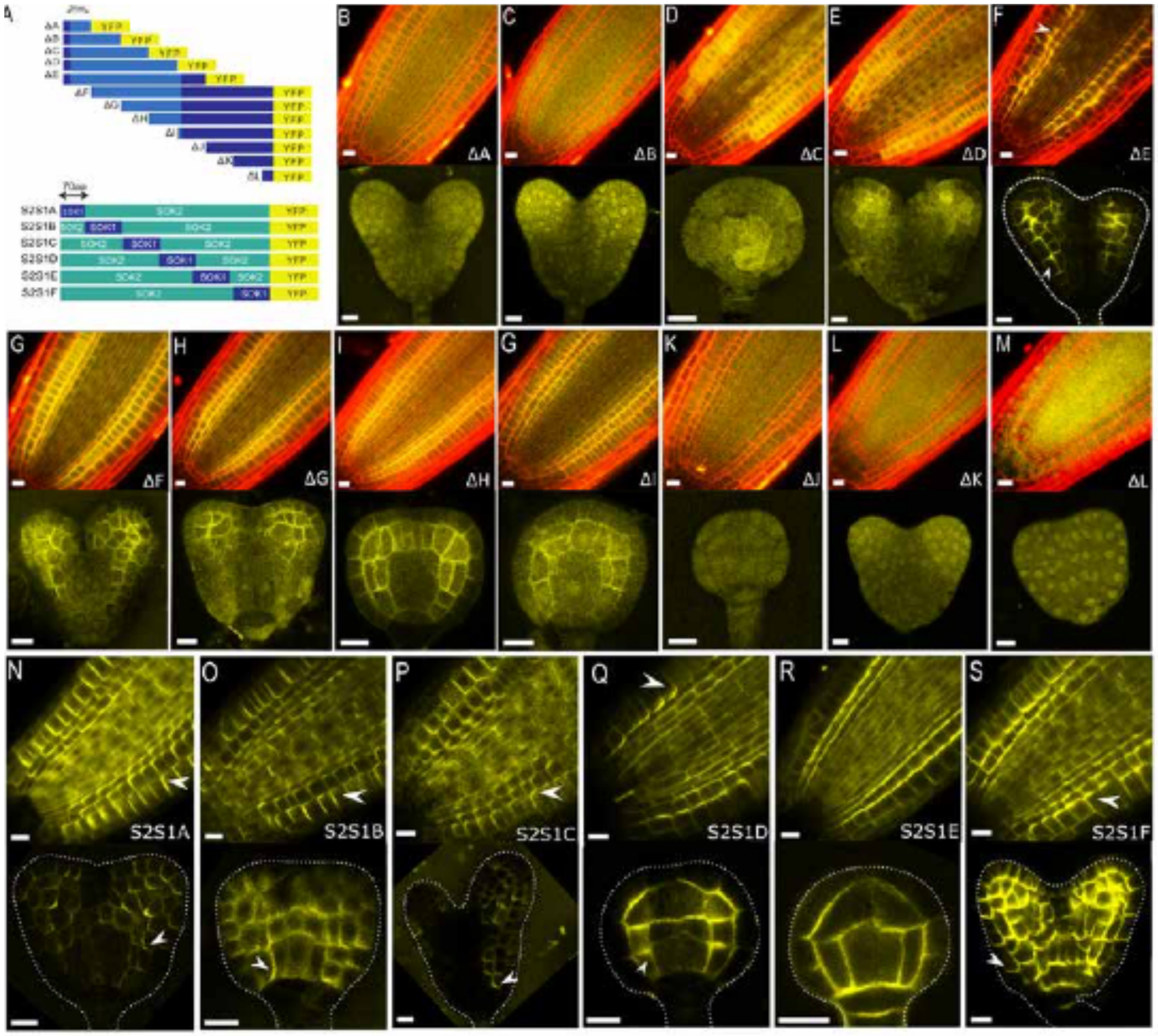


**Extended Figure 8: Details of SOK1 deletions and SOK1/2 swaps**
(a) Schematic of SOK1 domain deletion constructs expressed as YFP-fusions driven from the RPS5A promoter and SOK2-SOK1 domain swaps, also expressed as YFP fusions driven from the RPS5A promoter. (b-m) Complete set of SOK1-YFP domain deletion accumulation in root tips (upper panels, counterstained with Propidium Iodide – red) and heart stage embryos (lower panels). (n-s) Complete set of SOK1-SOK2-YFP domain swap accumulation in root tips (upper panels) and embryos (lower panels). Note that regions in (f,n-s) marked by arrows denote the regions for focused membrane localization and polarity defined by deletions and swaps. Bars 10 µm.

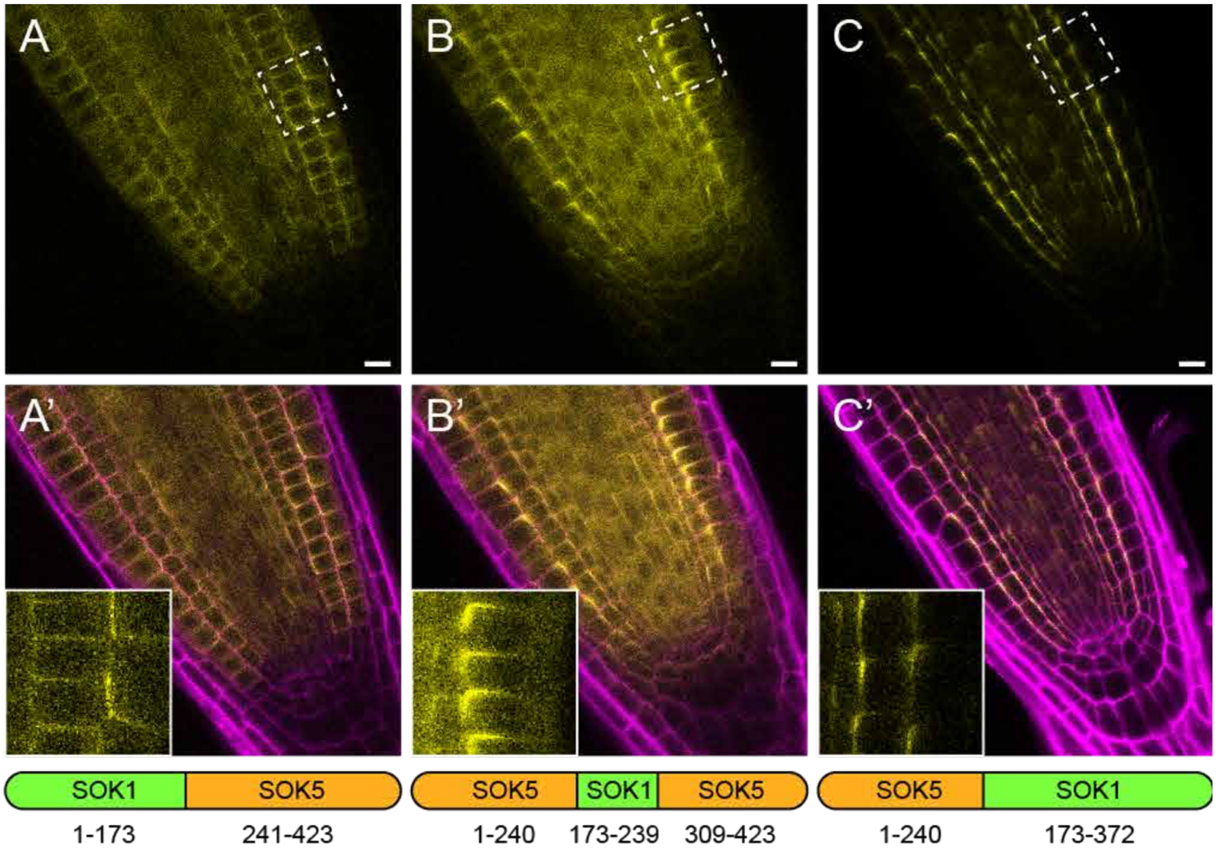


**Extended Figure 9: SOK1-SOK5 domain swaps**
Fluorescence of SOK1/SOK5 chimaeras, expressed as YFP fusions from the RPS5A promoter in root tips (a-c), and counterstained with Propidium Idodide (a'-c' – magenta). Insets in (a'-c') show protein localization in individual cells at high magnification (corresponding to dashed boxes in (a-c). Arrangement of SOK1 and SOK5 fragments in chimaeras are indicated underneath (a'-c'). Bars 10 µm.

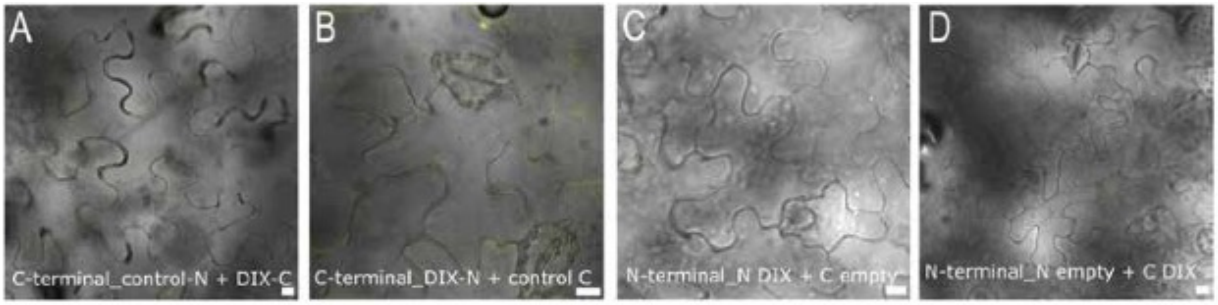


**Extended Figure 10: Controls for Bimolecular Fluorescence Complementation**
(a-d) Negative BiFC controls (single SOK1 DIX-cYFP or nYFP fusions with empty vector partner) in *Nicotiana benthamiana* epidermis. Bars 10 µm.

**Extended Table 1. List of treatments on SOK-YFP lines**

| treatment | conc (μM) | time |
|---|---|---|
| IAA | 1 | 1h |
| | | 4h |
| | | 6h |
| | | o.n. |
| | 10 | 6h |
| 2,4-D | 1 | o.n. |
| benzyladenine | 1 | o.n. |
| trans-zeatin | 1 | 1h |
| | | o.n. |
| epi-brassinolide | 1 | 1h |
| | | o.n. |
| GA | 1 | o.n. |
| NPA | 5 | 30min |
| | | 1-2h |
| | | 5h |
| | 10 | 6h |
| BFA | 42 | 30min |
| | | 1h |
| | | 6h |
| | | o.n. |
| wortmannin | 30 | 1h |
| | | o.n. |
| | 50 | 6h |
| concanamycinA | 0.8 | 1h |
| | | 6h |
| leptomycinB | 0.03 | 1.1h |
| | | 6h |
| | | o.n. |
| monensin | 10 | 6h |
| | | o.n. |
| cytochalasine B | 50 | 1h |
| | | 6h |
| | | o.n. |
| latrinculin B | 1 | 1.3h |
| | | 15min-2h |
| | | 6h |
| | | o.n. |
| jasplakinolide | 5 | 1h |
| | | o.n. |
| oryzalin | 5 | 1h |
| | | 6h |
| taxol | 100 | o.n. |
| isoxaben | 0.1 | 1 |
| | 1 | 1.5h |
| | 10 | 1.5h |
| | | 6h |
| MG-132 | 50 | 30min |
| | | 2h |
| | | 6h |
| | | o.n. |
| cycloheximide | 50 | 30min |
| | | 6h |

| | | |
|---|---|---|
| cycloheximide + MG-132 | 50, 50 | 1h |
| NPA + oryzalin | 5, 5 | 6h |
| temperature (37 degree) | | 1h |
| temperature (4 degree) | | 1h |
| gravity (90 degree) | | 1.5h |
| gravity (180 degree) | | 1.5h |
| dark | | 1d |

### Extended Table 2. Primers

TableS2. List of the primers used for cloning. All primer sequences are from 5' to 3'. LIC adapters are shown in red, Myr sequences are shown in blue, YFP sequences are shown in orange. Primers used multiple times are marked with the same Roman character.

| Construct | F/R | Primer sequence | Purpose | Used multiple times |
|---|---|---|---|---|
| SOK1-GFP | F | TAGTTGGAATGGGTTCGAACGTTCCGTGGTGAATCAATG | lic-pSOK1 | |
| | R | TTATGGAGTTGGGTTCGAACTCTCTTTCTTTTTGTTTTGGTCT | pSOK1-lic | |
| SOK2-GFP | F | TAGTTGGAATGGGTTCGAATGCCGATTCGATCTCGATACAGAG | lic-pSOK2 | A |
| | R | TTATGGAGTTGGGTTCGAACTTTTGTGTCTATAATGTCCGG | pSOK2-lic | |
| SOK3-GFP | F | TAGTTGGAATGGGTTCGAACGTTTATGGACTTACATTTCACTTAAGCATC | lic-pSOK3 | |
| | R | TTATGGAGTTGGGTTCGAAGTTTTTATCTCGGCGACCTAATTGG | pSOK3-lic | |
| SOK4-GFP | F | TAGTTGGAATGGGTTCGAACTCTCTTTCTGCTTCAGCTGAGTGAGATAGAG | lic-pSOK4 | B |
| | R | TTATGGAGTTGGGTTCGAATAATGTTGTTCGGTGTATTTTGGAGATTTG | pSOK4-lic | |
| SOK5-GFP | F | TAGTTGGAATGGGTTCGAAACTTAACGATCAAGAATTTAAGATAAGTCG | lic-pSOK5 | |
| | R | TTATGGAGTTGGGTTCGAACCTTTTTGTTTCTTTGCTTATGGAAAAGTG | pSOK5-lic | |
| pSOK1-SOK1-YFP | F | TAGTTGGAATGGGTTCGAATGAGTCGTTCCGTGGTGAATCAATG | lic-SOK1 | |
| | R | TTATGGAGTTGGGTTCGAACTCACTCTTTGAGAGTAGTCGTCAATACACG | SOK1-lic | |
| pSOK1-SOK1-tdT | F | TAGTTGGAATGGGTTCGAATGAGTCGTTCCGTGGTGAATCAATG | lic-SOK1 | |
| | R | TTATGGAGTTGGGTTCGAACTCACTCTTTGAGAGTAGTCGTCAATACACG | SOK1-lic | |
| pSOK2-SOK2-YFP | F | TAGTTGGAATGGGTTCGAATGCCGATTCGATCTCGATACAGAG | lic-SOK2 | A |
| | R | TTATGGAGTTGGGTTCGAACTTTCTTGATTTGCTTCGATGATGACATTAAACACG | SOK2-lic | |
| pSOK3-SOK3-YFP | F | TAGTTGGAATGGGTTCGAATGTGTGAGATTCATTTATCACCATACTTCAG | lic-SOK3 | |
| | R | TTATGGAGTTGGGTTCGAACAGGCTCTTCTGAAGGCACTTTCGA | SOK3-lic | |
| pSOK4-SOK4-YFP | F | TAGTTGGAATGGGTTCGAACTCTCTTTCTGCTTCAGCTGAGTGAGATAGAG | lic-SOK4 | B |
| | R | TTATGGAGTTGGGTTCGAACTGGTTGTCCAACCACAGAATTTGG | SOK4-lic | |
| pSOK5-SOK5-YFP | F | TAGTTGGAATGGGTTCGAAATCTAAAGTGTTATGTGCATACACACC | lic-SOK5 | |
| | R | TTATGGAGTTGGGTTCGAACCTGCTGCCCACCATTGTTCCT | SOK5-lic | |
| pRPS5A-SOK1 | F | TAGTTGGAATAGGTTCATGGAAAGTAATGGTGGAGGAGG | lic-SOK1 | |
| | R | AGTATGGAGTTGGGTTCTCACTCTTTGAGAGTAGTCGTCA | SOK1-lic | |
| pRPS5A-SOK1-YFP | F | TAGTTGGAATAGGTTCATGGAAAGTAATGGTGGAGGAGG | lic-SOK1 | C |
| | F | CGTGTATTGACGACTACTCTCAAAGAGATGACTAGTAAGGGCGAGGAGC | SOK1-YFP | D |
| | R | AGTATGGAGTTGGGTTCTTACTTGTACAGCTCGTCCATGCC | YFP-lic | E |
| pRPS5A-SOK2-YFP | F | TAGTTGGAATAGGTTCATGGAAGCTGTAAGATGCAGAAGA | lic-SOK2 | F |
| | F | CATCATCGAAGCAAATCAAGAAAATGACTAGTAAGGGCGAGGAGC | SOK2-YFP | G |
| | R | AGTATGGAGTTGGGTTCTTACTTGTACAGCTCGTCCATGCC | YFP-lic | E |
| Myr-SOK1 | F | TAGTTGGAATAGGTTCATGGGAGGATGCTTCTCTAAGAAGGAAAGTAATGGTGGAGGAGGAGAAG | lic-Myr-SOK1 | |
| SOK1-Myr | R | AGTATGGAGTTGGGTTCTTACTTCTTAGAGAAGCATCCTCCCTTGTACAGCTCGTCCATGCC | Myr-YFP | |
| deletion ΔA | F | CGGCTAAAACCACGTATAAACCAGGGAACCTGTTAAACC | pRPS5A | H |
| | R | GCCCTTACTAGTCATTAACCATTTCTTCAC | SOK1-YFP | |
| | F | GTGAAGAAATGGTTAATGACTAGTAAGGGC | SOK1-YFP | |
| | R | GCGGGACTCTAATCATAAAAACCCATCTCATAAATAACGTC | tNOS after YFP seq | I |
| | F | TAGTTGGAATAGGTTCATGGAAAGTAATGGTGGAGGAGG | lic-SOK1 | C |
| | R | AGTATGGAGTTGGGTTCTTACTTGTACAGCTCGTCCATGCC | YFP-lic | E |
| deletion ΔB | F | CGGCTAAAACCACGTATAAACCAGGGAACCTGTTAAACC | pRPS5A | H |
| | R | GCCCTTACTAGTCATTAGAAGAATCTCAGA | SOK1-YFP | |
| | F | TCTGAGATTCTTCTAATGACTAGTAAGGGC | SOK1-YFP | |
| | R | GCGGGACTCTAATCATAAAAACCCATCTCATAAATAACGTC | tNOS after YFP seq | I |
| | F | TAGTTGGAATAGGTTCATGGAAAGTAATGGTGGAGGAGG | lic-SOK1 | C |
| | R | AGTATGGAGTTGGGTTCTTACTTGTACAGCTCGTCCATGCC | YFP-lic | E |
| deletion ΔC | F | CGGCTAAAACCACGTATAAACCAGGGAACCTGTTAAACC | pRPS5A | H |
| | R | GCCCTTACTAGTCATCGATCTCTGTGAGCA | SOK1-YFP | |
| | F | TGCTCACAGAGATCGATGACTAGTAAGGGC | SOK1-YFP | |
| | R | GCGGGACTCTAATCATAAAAACCCATCTCATAAATAACGTC | tNOS after YFP seq | I |
| | F | TAGTTGGAATAGGTTCATGGAAAGTAATGGTGGAGGAGG | lic-SOK1 | C |
| | R | AGTATGGAGTTGGGTTCTTACTTGTACAGCTCGTCCATGCC | YFP-lic | E |
| deletion ΔD | F | CGGCTAAAACCACGTATAAACCAGGGAACCTGTTAAACC | pRPS5A | H |
| | R | CCTTACTAGTCATACTTGGTCGACCCGA | SOK1-YFP | |
| | F | TCGGGTCGACCAAGTATGACTAGTAAGG | SOK1-YFP | |
| | R | GCGGGACTCTAATCATAAAAACCCATCTCATAAATAACGTC | tNOS after YFP seq | I |
| | F | TAGTTGGAATAGGTTCATGGAAAGTAATGGTGGAGGAGG | lic-SOK1 | C |
| | R | AGTATGGAGTTGGGTTCTTACTTGTACAGCTCGTCCATGCC | YFP-lic | E |
| deletion ΔE | F | CGGCTAAAACCACGTATAAACCAGGGAACCTGTTAAACC | pRPS5A | H |
| | R | GCCCTTACTAGTCATCCCAGACCTAGACTT | SOK1-YFP | |
| | F | AAGTCTAGGTCTGGGATGACTAGTAAGGGC | SOK1-YFP | |
| | R | GCGGGACTCTAATCATAAAAACCCATCTCATAAATAACGTC | tNOS after YFP seq | I |
| | F | TAGTTGGAATAGGTTCATGGAAAGTAATGGTGGAGGAGG | lic-SOK1 | C |
| | R | AGTATGGAGTTGGGTTCTTACTTGTACAGCTCGTCCATGCC | YFP-lic | E |
| Template for ΔF-ΔL | F | CGGCTAAAACCACGTATAAACCAGGGAACCTGTTAAACC | pRPS5A | H |
| | R | GCTCCTCGCCCTTACTAGTCATCTCTTTGAGAGTAGTCGTCAATACACG | SOK1-YFP | J |
| | F | CGTGTATTGACGACTACTCTCAAAGAGATGACTAGTAAGGGCGAGGAGC | SOK1-YFP | D |
| | R | GCGGGACTCTAATCATAAAAACCCATCTCATAAATAACGTC | tNOS after YFP seq | I |
| deletion ΔF | F | TAGTTGGAATAGGTTCATGGCAGACGCAC | lic-SOK1 | |
| | R | AGTATGGAGTTGGGTTCTTACTTGTACAGCTCGTCCATGCC | YFP-lic | E |
| deletion ΔG | F | TAGTTGGAATAGGTTCATGAGCTCTCCAAAG | lic-SOK1 | |
| | R | AGTATGGAGTTGGGTTCTTACTTGTACAGCTCGTCCATGCC | YFP-lic | E |
| deletion ΔH | F | TAGTTGGAATAGGTTCATGACGGCAACAAC | lic-SOK1 | |
| | R | AGTATGGAGTTGGGTTCTTACTTGTACAGCTCGTCCATGCC | YFP-lic | E |
| deletion ΔI | F | TAGTTGGAATAGGTTCATGGTATCGTCTAC | lic-SOK1 | |
| | R | AGTATGGAGTTGGGTTCTTACTTGTACAGCTCGTCCATGCC | YFP-lic | E |
| deletion ΔJ | F | TAGTTGGAATAGGTTCATGGCTTTTGGTCC | lic-SOK1 | |
| | R | AGTATGGAGTTGGGTTCTTACTTGTACAGCTCGTCCATGCC | YFP-lic | E |
| deletion ΔK | F | TAGTTGGAATAGGTTCATGCCTTCCATGGC | lic-SOK1 | |
| | R | AGTATGGAGTTGGGTTCTTACTTGTACAGCTCGTCCATGCC | YFP-lic | E |
| deletion ΔL | F | TAGTTGGAATAGGTTCATGAGAAACATTCC | lic-SOK1 | |
| | R | AGTATGGAGTTGGGTTCTTACTTGTACAGCTCGTCCATGCC | YFP-lic | E |
| swap S2S1A | F | CGGCTAAAACCACGTATAAACCAGGGAACCTGTTAAACC | pRPS5A | H |
| | R | TGTGAGATAGTAAACAACTTGGACTCTTCGTACTTCTCCTCCTCCACCATTACTTTCCAT | SOK1-SOK2 | |
| | F | ATGGAAAGTAATGGTGGAGGAGGAGAAGTACGAAGAGTCCAAGTTGTTTACTATCTCAC | SOK1-SOK2 | |
| | R | GGTGAACAGCTCCTCGCCCTTACTAGTCATTTTCTTGATTTGCTTCGATGATGACATTAA | SOK2-YFP | K |
| | F | TTAATGTCATCATCGAAGCAAATCAAGAAAATGACTAGTAAGGGCGAGGAGCTGTTCACC | SOK2-YFP | G |
| | R | GCGGGACTCTAATCATAAAAACCCATCTCATAAATAACGTC | tNOS after YFP seq | I |
| | F | TAGTTGGAATAGGTTCATGGAAAGTAATGGTGGAGGAGG | lic-SOK1 | C |
| | R | AGTATGGAGTTGGGTTCTTACTTGTACAGCTCGTCCATGCC | YFP-lic | E |
| swap S2S1B | F | CGGCTAAAACCACGTATAAACCAGGGAACCTGTTAAACC | pRPS5A | H |
| | R | GCTAAGGAAATAAACTAAATTCACTCTTCTGAAAATGGGTTTCTTTGTCTTGACCTCTTC | SOK2-SOK1 | |

| | | | | |
|---|---|---|---|---|
| | F | GAAGAGGTCAAGACAAAGAAACCCATTTTCAGAAGAGTGAATTTAGTTTATTTCCTTAGC | SOK2-SOK1 | |
| | R | TGGTCTGTTCACATGTACTTCTTGAAATTTTAGAAGAATCTCAGATCCTTTGAGGACATA | SOK1-SOK2 | |
| | F | TATGTCCTCAAAGGATCTGAGATTCTTCTAAAATTTCAAGAAGTACATGTGAACAGACCA | SOK1-SOK2 | |
| | R | GGTGAACAGCTCCTCGCCCTTACTAGTCATTTTCTTGATTTGCTTCGATGATGACATTAA | SOK2-YFP | K |
| | F | TTAATGTCATCATCGAAGCAAATCAAGAAAATGACTAGTAAGGGCGAGGAGCTGTTCACC | SOK2-YFP | G |
| | R | GCGGGACTCTAATCATAAAAACCCATCTCATAAATAACGTC | tNOS after YFP seq | I |
| | F | TAGTTGGAATAGGTTCATGGAAGCTGTAAGATGCAGAAGAGG | lic-SOK2 | F |
| | R | AGTATGGAGTTGGGTTCTTACTTGTACAGCTCGTCCATGCC | YFP-lic | E |
| swap S2S1C | F | CGGCTAAAACCACGTATAAACCAGGGAACCTGTTAAACC | pRPS5A | H |
| | R | CACATTAGGATAATCCTCCTTTGGAGAGCTGTCGGTGATTTCAGATCCTTTCAAGACATA | SOK2-SOK1 | |
| | F | TATGTCTTGAAAGGATCTGAAATCACCGACAGCTCTCCAAAGGAGGATTATCCTAATGTG | SOK2-SOK1 | |
| | R | TACTTGCAAATCTTGCTCTGTCTTGGTCAACTTAAGAACGAAACCTTCTTCGTTCGTGGT | SOK1-SOK2 | |
| | F | ACCACGAACGAAGAAGGTTTCGTTCTTAAGTTGACCAAGACAGAGCAAGATTTGCAAGTA | SOK1-SOK2 | |
| | R | GGTGAACAGCTCCTCGCCCTTACTAGTCATTTTCTTGATTTGCTTCGATGATGACATTAA | SOK2-YFP | K |
| | F | TTAATGTCATCATCGAAGCAAATCAAGAAAATGACTAGTAAGGGCGAGGAGCTGTTCACC | SOK2-YFP | G |
| | R | GCGGGACTCTAATCATAAAAACCCATCTCATAAATAACGTC | tNOS after YFP seq | I |
| | F | TAGTTGGAATAGGTTCATGGAAGCTGTAAGATGCAGAAGAGG | lic-SOK2 | F |
| | R | AGTATGGAGTTGGGTTCTTACTTGTACAGCTCGTCCATGCC | YFP-lic | E |
| swap S2S1D | F | CGGCTAAAACCACGTATAAACCAGGGAACCTGTTAAACC | pRPS5A | H |
| | R | CTGGCCGGAGACTGTTTTCGGATCTTGTTTGTTCGGTTTTTGTTCGGTGGATTCCATTGT | SOK2-SOK1 | |
| | F | ACAATGGAATCCACCGAACAAAAACCGAACAAACAAGATCCGAAAACAGTCTCCGGCCAG | SOK2-SOK1 | |
| | R | TCTCGGATTCATCACACTCGGTGCGTAGTAGTTTGTGTCCAAACCACCACACTTCATCAA | SOK1-SOK2 | |
| | F | TTGATGAAGTGTGGTGGTTTGGACACAAACTACTACGCACCGAGTGTGATGAATCCGAGA | SOK1-SOK2 | |
| | R | GGTGAACAGCTCCTCGCCCTTACTAGTCATTTTCTTGATTTGCTTCGATGATGACATTAA | SOK2-YFP | K |
| | F | TTAATGTCATCATCGAAGCAAATCAAGAAAATGACTAGTAAGGGCGAGGAGCTGTTCACC | SOK2-YFP | G |
| | R | GCGGGACTCTAATCATAAAAACCCATCTCATAAATAACGTC | tNOS after YFP seq | I |
| | F | TAGTTGGAATAGGTTCATGGAAGCTGTAAGATGCAGAAGAGG | lic-SOK2 | F |
| | R | AGTATGGAGTTGGGTTCTTACTTGTACAGCTCGTCCATGCC | YFP-lic | E |
| swap S2S1E | F | CGGCTAAAACCACGTATAAACCAGGGAACCTGTTAAACC | pRPS5A | H |
| | R | AGACTTATTCAAAGGCACCAATACGGCGTCCTTGGTGGCGATGTGACCGCAAGAGATCAT | SOK2-SOK1 | |
| | F | ATGATCTCTTGCGGTCACATCGCCACCAAGGACGCCGTATTGGTGCCTTTGAATAAGTCT | SOK2-SOK1 | |
| | R | TTCAGCCGTGACTCGTTCCTGACTCACCGACGAGCATAATGGAGCCATGGAAGGCTTGCT | SOK1-SOK2 | |
| | F | AGCAAGCCTTCCATGGCTCCATTATGCTCGTCGGTGAGTCAGGAACGAGTCACGGCTGAA | SOK1-SOK2 | |
| | R | GGTGAACAGCTCCTCGCCCTTACTAGTCATTTTCTTGATTTGCTTCGATGATGACATTAA | SOK2-YFP | K |
| | F | TTAATGTCATCATCGAAGCAAATCAAGAAAATGACTAGTAAGGGCGAGGAGCTGTTCACC | SOK2-YFP | G |
| | R | GCGGGACTCTAATCATAAAAACCCATCTCATAAATAACGTC | tNOS after YFP seq | I |
| | F | TAGTTGGAATAGGTTCATGGAAGCTGTAAGATGCAGAAGAGG | lic-SOK2 | F |
| | R | AGTATGGAGTTGGGTTCTTACTTGTACAGCTCGTCCATGCC | YFP-lic | E |
| swap S2S1F | F | CGGCTAAAACCACGTATAAACCAGGGAACCTGTTAAACC | pRPS5A | H |
| | R | CTTCTCAGGTTTGAACAACTTCCCACATTGCTCAACGATGCTTCCGCTGAAATACTCTTT | SOK2-SOK1 | |
| | F | AAAGAGTATTTCAGCGGAAGCATCGTTGAGCAATGTGGGAAGTTGTTCAAACCTGAGAAG | SOK2-SOK1 | |
| | R | GGTGAACAGCTCCTCGCCCTTACTAGTCATCTCTTTGAGAGTAGTCGTCAATACACGTTG | SOK1-YFP | J |
| | F | CAACGTGTATTGACGACTACTCTCAAAGAGATGACTAGTAAGGGCGAGGAGCTGTTCACC | SOK1-YFP | D |
| | R | GCGGGACTCTAATCATAAAAACCCATCTCATAAATAACGTC | tNOS after YFP seq | I |
| | F | TAGTTGGAATAGGTTCATGGAAGCTGTAAGATGCAGAAGAGG | lic-SOK2 | F |
| | R | AGTATGGAGTTGGGTTCTTACTTGTACAGCTCGTCCATGCC | YFP-lic | E |
| UAS::SOK1-YFP | F | TAGTTGAATAGGTTCATGGAAAGTAATGGTGGAGG | LIC adaptor -SOK1 | |
| | R | AGTATGGAGTTGGGTTCTTACTTGTACAGCTCGTCC | YFP-LIC adaptor | |
| 35S::SOK1DIX-YFP/CFP FLIM | F | TAGTTGGAATAGGTTCATGGAAAGTAATGGTGGAG | LIC adaptor -SOK1 DIX | |
| | R | AGTATGGAGTTGGGTTCCTCCTTTGGAGAGCTTAG | SOK1 DIX-LIC Adaptor | |
| 35S::SOK1DIX-n/cYFP BiFC | F | TAGTTGGAATAGGTTCATGGAAAGTAATGGTGGAG | LIC adaptor -SOK1 DIX | |
| | R | AGTATGGAGTTGGGTTCGACTCCTTTGGAGAGCTTAG | SOK1 DIX-LIC Adaptor | |
| 35S::n/cYFP-SOK1DIX BiFC | F | TAGTTGGAATAGGTTCATGGAAAGTAATGGTGGAG | LIC adaptor -SOK1 DIX | |
| | R | AGTATGGAGTTGGGTTCTTACTCCTTTGGAGAGCTTAGAA | SOK1 DIX-LIC Adaptor | |
| swap A-E B | F | TAGTTGGAATAGGTTCATGGAAAGTAATGGTGGAGGAGG | lic-SOK1 | C |
| | F | GAACGAAGAAGGTTTCGTTCTTAGCAGAGACGAGATATCTCC | SOK1-SOK5 | |
| | R | GGAGATATCTCGTCTCTGCTAAGAACGAAACCTTCTTCGTTC | SOK5-SOK1 | |
| | R | AGTATGGAGTTGGGTTCTTACTTGTACAGCTCGTCCATGCC | YFP-lic | E |
| swap A-E C | F | TAGTTGGAATAGGTTCATGAGTTCAAGAGTGTTCAGAGC | LIC-SOK5 | L |
| | F | GAATCAGAGTACAGAGCTGCAGAGCTGAAGAAACAAGATCCGAAAACAG | SOK5-SOK1 | |
| | F | GTGGTGGTTTGGACACAAACGAATGTGGGCCTGTTTTGTTG | SOK1-SOK5 | |
| | R | CTGTTTTCGGATCTTGTTTCTTCAGCTCTGTACTCTGATTC | SOK5-SOK1 | |
| | R | CAACAAAACAGGCCCACATTCGTTTGTGTCCAAACCACCAC | SOK1-SOK5 | |
| | R | AGTATGGAGTTGGGTTCTTACTTGTACAGCTCGTCCATGCC | YFP-lic | E |
| swap A-E D | F | TAGTTGGAATAGGTTCATGAGTTCAAGAGTGTTCAGAGC | LIC-SOK5 | L |
| | F | GAATCAGAGTACAGAGCTGAAGAAACAAGATCCGAAAACAG | SOK5-SOK1 | |
| | R | CTGTTTTCGGATCTTGTTTCTTCAGCTCTGTACTCTGATTC | SOK1-SOK5 | |
| | R | AGTATGGAGTTGGGTTCTTACTTGTACAGCTCGTCCATGCC | YFP-lic | E |

**Supplemental Movie 1:**
Localization of SOK1-YFP in heart stage embryo. Embryo was stained with Renaissance RS2200 (white), observed by confocal, and 3D reconstructed.

**Supplemental Movie 2:**
A time-lapse series of SOK1-YFP during lateral root initiation. Root was stained with Propidium Iodide and observed by confocal. Time (hours) is indicated.

**Supplemental Movie 3:**
A time-lapse series of SOK1-YFP during root growth. Root was stained with Propidium Iodide and observed by vertical confocal. Time (hours) is indicated.
